# Biomarker evaluation of plasma microRNA-122, high-mobility group box-1 and keratin-18 in acute gallstone disease

**DOI:** 10.1101/189589

**Authors:** Francesca Th’ng, Bastiaan Vliegenthart, Jonathan D Lea, Daniel J. Antoine, James W Dear, Damian J. Mole

## Abstract

**Background:** A biomarker that stratifies patients with complications of gallstone disease from the denominator pool of people with acute biliary-type symptoms is needed. Circulating microRNA-122 (miRNA-122), high-mobility group box1 (HMGB1), full-length keratin-18 (flk-18) and caspase-cleaved keratin-18 (cck-18) are established hepatocyte injury biomarkers. The aim of this study was to evaluate the discriminatory power of these biomarkers in plasma to identify potential biliary complications that may require acute intervention.

**Method:** An observational biomarker cohort study was carried out in a University teaching hospital for 12 months beginning 3^rd^ September 2014. Blood samples were collected from adults referred with acute biliary-type symptoms. miRNA-122 was measured by quantitative real-time polymerase chain reaction, and HMGB1, cck-18 and flk-18 by ELISA.

**Results:** 300 patients were screened and 289 patients were included. Plasma miRNA-122, cck-18 and flk-18 concentrations were increased in patients with gallstones compared to those without (*miRNA-122*: median: 2.89 × 10^4^ copies/ml vs. 0.95 × 10^4^ copies/ml [P<0.001]; *cck-18:* 121.9 U/L vs. 104.6 U/L [P = 0.041]; *flk-18*: 252.4 U/L vs. 151.8 U/L [P<0.001]). Uncomplicated gallstone disease was associated with higher miRNA-122 and cck-18 concentrations than complicated disease (*miRNA-122*: 5.72 × 10^4^ copies/ml vs. 2.26x10^4^ copies/ml [P=0.022]; *cck-18*: 139.7 U/L vs. 111.4 U/L [P=0.049]). There was no significant difference in HMGB1 concentration between patients with and without gallstones [P=0.480]. Separation between groups for all biomarkers was modest.

**Conclusion:** microRNA-122 and keratin-18 plasma concentrations are elevated in patients with gallstones. However, these biomarkers were not sufficiently discriminatory to be progressed as clinically useful biomarkers in this context.

## INTRODUCTION

Gallstone disease is one of the most common reasons for persons to present to hospital with gastrointestinal disease in Western countries^1^. In the United Kingdom, there are approximately 330 people per 100,000 adult population admitted with an acute biliary disease annually^2^. A point-of-care diagnostic test that can stratify patients with acute biliary-type symptoms according to their need for further investigation or intervention will be advantageous to patients and has the potential to reduce healthcare expenditure. Individuals with uncomplicated biliary colic and aseptic choledocholithiasis may be safely discharged without the need for inpatient management and can be managed in an outpatient or day-case setting. Conversely, patients with complications, for example cholecystitis, cholangitis and gallstone pancreatitis, are best managed as inpatients according to current national standards.

Standard available clinico-pathological data for patients presenting acutely with gallstones in the emergency setting include clinical history and examination, routine laboratory results, and ultrasound (USS) or magnetic resonance imaging (MRI) findings^3^. Currently, this set of investigations cannot always identify whether patients have complications of gallstones that necessitate admission. Therefore, a biomarker panel that could be used as an adjunct to existing investigations to guide treatment decisions would be extremely advantageous.

microRNA-122 (miRNA-122), high mobility group box 1 (HMGB1), full-length keratin-18 (flk-18) and caspase-cleaved keratin-18 (cck-18) have been identified as early acute hepatic injury biomarkers at presentation to hospital, particularly in the context of paracetamol overdose^4,5^. miRNAs are single-stranded, small non-protein-coding ribonucleic acids (RNAs). miRNA-122 is expressed near exclusively in the hepatocyte and provides enhanced hepatic specificity over all current liver injury biomarkers^6^. HMGB1 is a nuclear binding protein with pro-inflammatory activity^7^ that is released by immune and necrotic cells^7,8^. Keratins are part of the intermediate filament system and they play an important role in cellular mechanisms. They are a hallmark of epithelial cells including hepatocytes and biliary cells^9,10^. During apoptosis, full-length keratin-18 (flk-18) is cleaved by caspases to produce the fragment caspase-cleaved keratin-18 (cck-18)^11^. flk-18 is released into blood during necrosis and cck-18 during apoptosis^12^. Studies in pre-clinical models have demonstrated increased circulating miRNA-122, HMGB1 and keratin-18 following bile duct ligation^13^. In humans with cholecystitis, keratin-18 has been reported to be elevated in both bile and serum samples, when compared to a control group of healthy volunteers^14^. Shifeng *et al.*^6^ reported that circulating miRNA-122 has a high diagnostic value with regard to distinguishing patients with biliary calculi from normal controls. In that study, the area under the receiver operator curve (ROC-AUC) was 0.93, which suggests high accuracy. However, the controls were healthy subjects, whereas in clinical practice, the relevant comparison group will be patients with other aetiologies of abdominal pain. In the present study, we recruited a cohort of patients with acute abdominal pain.

The aim of this study was to evaluate whether circulating hepatic injury biomarkers discriminate between people with and without gallstone disease and uncomplicated from complicated gallstone disease during the first 24 hours of hospital admission.

## METHODS

### Ethics

This study was assessed by South East Scotland Research Ethics Service of National Health Service (NHS) Lothian and was given ethical approval under the terms of the Governance Arrangements for Research Ethics Committees (Harmonised Edition). Lothian R&D project number 2014/0224; REC number 14/EM/0211.

### Study Design and Setting

This study was an observational cohort biomarker study conducted in the Emergency Department and Emergency General Surgery Department at the Royal Infirmary of Edinburgh, NHS Lothian, U.K.

### Inclusion and Exclusion Criteria

All adult patients referred to the on-call General Surgeons with acute abdominal pain, irrespective of the characteristics of the pain, and with a differential diagnosis of gallstone disease upon index presentation were eligible for inclusion. Patients must have had presented to hospital within 24 hours prior to recruitment. Patients who were unable to give informed consent, or who refused to participate in the study, or who were younger than 16 years of age or transferred from a hospital outside the NHS regional trust (NHS Lothian) were excluded from the study.

### Study Population, sample size and data collection

This proof-of-concept research study was designed to inform a power calculation for a multi-centre qualification study, and therefore the sample size was pragmatic. This study had been approved to recruit a total of 300 patients over a period of two years.

Recruitment was for 52 weeks: 3rd September 2014 to 2nd September 2015. Potential participants were identified at the on-call General Surgery team’s daily handover. All patients recruited to the study had given informed consent. Clinical details, imaging reports, operation findings and pathology reports were obtained from hand written and electronic medical notes. Each patient was then followed up through the electronic record for a minimum of three months. Whole blood was sampled within 24 hours of hospital presentation into EDTA vaccutainer tubes (S-Monovette, Sarstedt), centrifuged at 1000xg for 15minutes at 4°C and the plasma layer retained. The supernatant was then separated into aliquots and stored at -80°C until analysis.

### Variables and Definitions

Clinical data included patient demographics, routine laboratory blood and microbiology results, imaging investigations, and/or information on surgical interventions and pathology results. The final diagnoses of the patients were determined by histopathology results, operative findings, imaging investigations and/or discharge scripts, following that hierarchy. Patients with a diagnosis of non-specific abdominal pain (NSAP) had normal investigation results and a definitive cause of their abdominal pain was not found. Patients with gallstone diseases were grouped into ‘Uncomplicated’ and ‘Complicated’ categories: patients with ‘uncomplicated’ disease had either biliary colic (cholelithiasis in gallbladder) or aseptic choledocholithiasis (choledocholithiasis without clinical, laboratory or microbiological evidence of sepsis); patients with ‘complicated’ gallstone disease included those with cholangitis (septic choledocholithiasis), cholecystitis or gallstone pancreatitis.

### Total RNA Extraction, Quantitative Real-Time Polymerase Chain Reaction (qRT-PCR) and enzyme-linked immunosorbent assay (ELISA) Analysis

Laboratory analyses of all biomarkers were carried out blinded to participant clinical data. Analysis of miRNA-122 was carried out in the Centre for Cardiovascular Sciences, University of Edinburgh); HMGB1, flk-18 and cck-18 laboratory analyses were carried out at the MRC Centre for Drug Safety Science, University of Liverpool.

miRNA was extracted from plasma using the miRNeasy Serum/Plasma Kit (Qiagen, Venlo, Netherlands) according to the manufacturer’s protocol^15,16^. Synthetic miR-39 (at 1.6 × 10^8^ copies/μL) was spiked in as an internal control. miRNAs were measured with Taqman-based qPCR. Small RNA elutes were reverse transcribed using specific stem-loop reverse-transcription RT primers (Applied Biosystems, Foster City, CA, USA) for each target miRNA species, following the manufacturer’s instructions. Specific stem loop rt primer targeting UGGAGUGUGACAAUGGUGUUUG and 5’ UCACCGGGUGUAAAUCAGCUUG 3’ was used. No template controls (NTCs) were included to test for miRNA contamination. The expression of miR-39 and miRNA-122 were analysed using the standard 2^-dct^ method^17^ normalized to the miR-39 spiked internal control.

Plasma HMGB1 and keratin-18 were determined by ELISA according to the manufacturer’s guidelines (Shino-Test/IBL International^18,19^ for HMGB1; and PEVIVA^20^ for cck-18 and flk-18) and our previously published protocols^5,21^.

### Statistical analysis

Continuous variables are presented as medians and interquartile ranges (IQR). Categorical variables are presented as frequencies. The Pearson Chi-squared test was used to examine the association between categorical variables. The Kolmogorov-Smirnov test was used to compare the distribution of non-parametric data against the normal distribution. The Kruskal-Wallis test was used to compare non-parametric variables followed by Dunn’s *post hoc* test for multiple comparisons. 95% confidence intervals (CI) were calculated for all normally distributed continuous variables. Positive predictive value (PPV) and negative predictive value (NPV) were utilized as the most relevant measures of clinical utility. Receiver-operating characteristic (ROC) curve analyses were plotted for the different patient groups. Areas under the ROC curves were calculated with 95% CI. Sensitivity, PPV and NPV were obtained at 90% specificity. All statistical tests were based on a two-sided *α*–value of 0.05. Statistical analysis was performed using IBM SPSS Statistics Version 19.0 (IBM Corp., Armonk, NY, USA), and G*Power3.1 (Universität Düsseldorf, Germany). Figures were designed, and correlation analyses were carried out using GraphPad Prism Version 6.0 (GraphPad Software, Inc., La Jolla, CA, USA).

## RESULTS

### Study Population

A total of 300 patients were screened for possible inclusion in the study. The study CONSORT flow-chart is presented in Figure 1. Eleven patients who had consented to participate in the study were excluded: two were transferred to a different hospital prior obtaining blood samples; one was under 16 years old; five were admitted more than 24 hours prior to recruitment; two patients’ blood samples were collected in the wrong sample tube; one blood sample tube did not have the required volume for biomarker testing. Of the 289 patients included in the study, as their final diagnosis, 47 (16.3%) patients had nonspecific abdominal pain (NSAP), 64 patients (22.1%) had uncomplicated gallstone disease, 115 patients (39.8%) had a complicated gallstone disease and 63 patients (21.8%) had other non-gallstone related disease.

**Figure 1:**
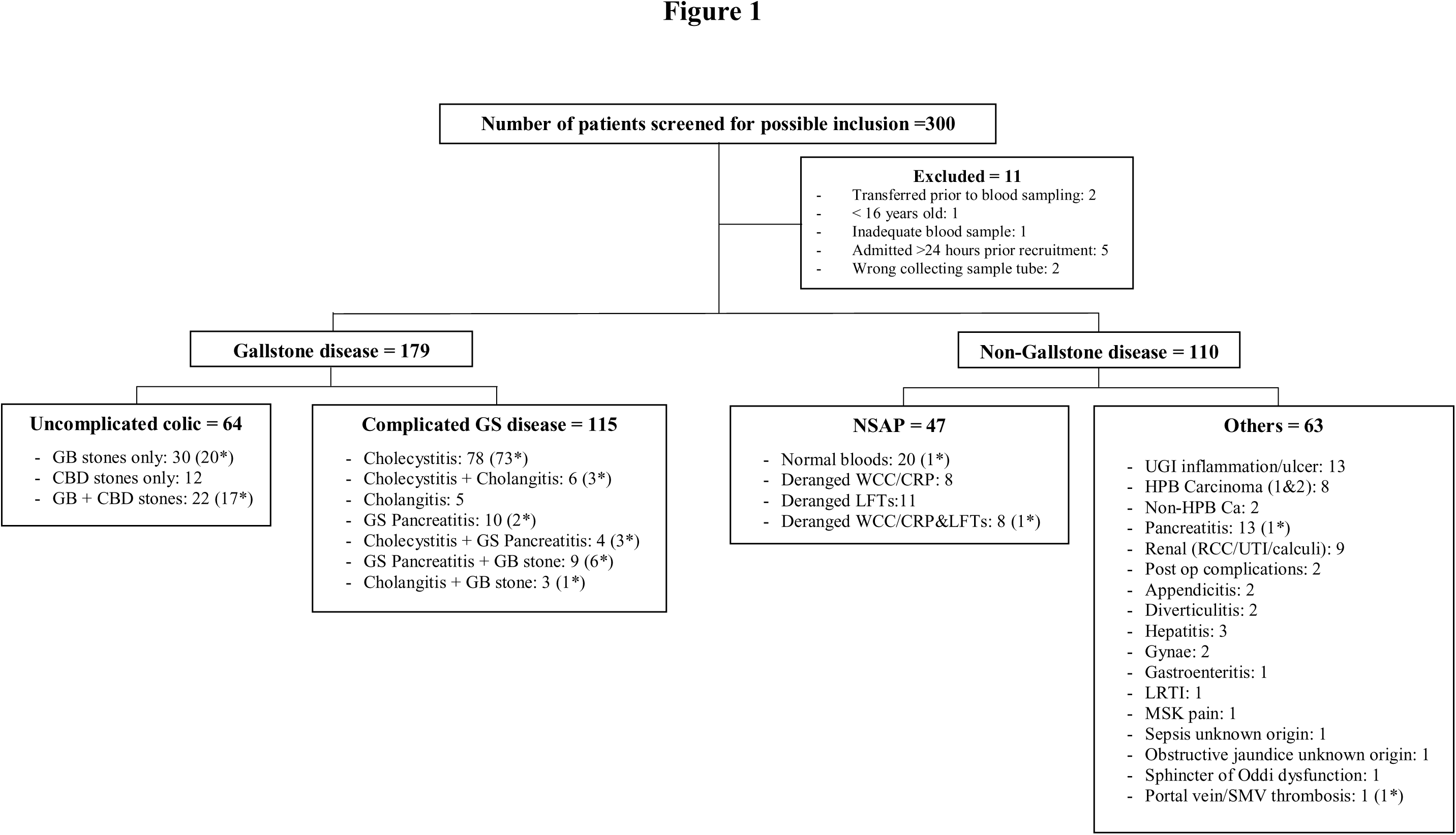
CONSORT diagram for the main study. The number of patients who underwent cholecystectomy within each group is illustrated in brackets with an asterisk). GS, gallstones; GB, gall bladder; CBD, common bile duct; NSAP, non-specific abdominal pain; UGI, upper gastrointestinal; HPB, hepatopancreaticobiliary; RCC, renal cell cancer; UTI, urinary tract infection; LRTI, lower respiratory tract infection; MSK, musculoskeletal; SMV, superior mesenteric vein; WCC, white blood cell count; CRP, C-reactive protein; LFT, liver function test.

The demographics of the study subjects are illustrated in Table 1. There were more females compared to males between the four patient groups (P<0.001). Patients with uncomplicated gallstone diseases (46.4 years [I29.8-61.8]) were significantly younger than patients with complicated diseases (56.6 years [44.6-68.5]; P=0.002).

**Table 1.** Demographics of the study cohort and routine blood test results.

|  | Overall | NSAP<sup>1</sup> | Uncomplicated<sup>2</sup> | Complicated<sup>3</sup> | Others | P-value |
| --- | --- | --- | --- | --- | --- | --- |
| N (%) | 289 (100) | 47 (16.3) | 64 (22.1) | 115 (39.8) | 63 (21.8) |  |
| Gender (n, (%)) |  |  |  |  |  |  |
| Male | 88 (30.4) | 3 (6.4) | 9 (14.1) | 53 (46.1) | 23 (36.5) | <0.001 |
| Female | 201 (69.6) | 44 (93.6) | 55 (85.9) | 62 (53.9) | 40 (63.5) |  |
| Age (Years) |  |  |  |  |  |  |
| Median | 51.3 | 41.3 | 46.4 | 56.6 | 52.2 | <0.001 |
| IQR | 37.5-67.4 | 26.7-57.3 | 29.8-61.8 | 44.6-68.5 | 39.3-67.5 |  |
| 95% CI | 49.3,53.5 | 37.9,49.4 | 42.7,51.7 | 53.1,59.4 | 48.2,56.6 |  |
| Underwent Operation (n, (%)) |  |  |  |  |  |  |
| Yes | 276 (44.6) | 2 (4.3) | 37 (57.8) | 88 (76.5) | 2 (3.2) | <0.001 |
| No | 160 (55.4) | 45 (95.7) | 27 (42.2) | 27 (23.5) | 61 (96.8) |  |
| Duration from index presentation to Operation (Days) |  |  |  |  |  |  |
| Median | 3.0 | 1.5 | 4.0 | 3.0 | 24.0 | 0.067 |
| IQR | 2.0-5.5 | 1.0- - | 2.0-7.5 | 2.0-4.8 | 5.0- - |  |
| 95% CI | 7.6,17.6 | -4.9,7.9 | 4.1,25.2 | 5.8,17.6 | -217.4,265.4 |  |
| Total bilirubin (umol/L) |  |  |  |  |  |  |
| Median | 14.0 | 11.0 | 22.5 | 17.0 | 11.0 | <0.001 |
| IQR | 8.0-31.5 | 7.0-15.0 | 10.3-46.8 | 9.0-35.0 | 7.0-33.0 |  |
| 95% CI | 24.3,32.7 | 10.2,14.2 | 25.9,44.1 | 23.3,35.6 | 19.8,44.6 |  |
| Alanine aminotransferase, ALT (U/L) |  |  |  |  |  |  |
| Median | 36.0 | 23.0 | 135.0 | 38.0 | 41.0 | <0.001 |
| IQR | 19.0-169.5 | 14.0-29.0 | 23.5-359.0 | 20.0-178.0 | 18.0-100.0 |  |
| 95% CI | 119.2,174.1 | 20.9,32.2 | 175.7,336.5 | 106.6,185.2 | 71.4,181.5 |  |
| Alkaline Phosphatases, AlkP (U/L) |  |  |  |  |  |  |
| Median | 96.0 | 76.0 | 139.5 | 94.0 | 94.0 | <0.001 |
| IQR | 71.0-172.5 | 60.0-103.0 | 86.0-264.0 | 70.0-158.0 | 74.0-238.0 |  |
| 95% CI | 137.9,172.0 | 75.2,101.3 | 153.1,223.0 | 121.7,170.5 | 135.8,238.7 |  |
| Gamma-glutamyl transpeptidase, GGT (U/L) |  |  |  |  |  |  |
| Median | 87.0 | 25.0 | 238.0 | 114.5 | 74.0 | <0.001 |
| IQR | 30.0-389.5 | 14.0-54.0 | 55.0-457.5 | 39.3-392.5 | 30.0-497.8 |  |
| 95% CI | 211.4,312.7 | 24.9,54.5 | 236.5,422.1 | 178.9,358.3 | 187.4,437.3 |  |

<sup>1</sup> Non-Specific Abdominal Pain<sup>2</sup> Biliary colic, Aseptic Choledocholithiasis<sup>3</sup> Cholecystitis, Cholangitis, Gallstone Pancreatitis<sup>4</sup> Pearson Chi-Square test (For 'Gender' and 'Underwent Operation')<sup>5</sup> Kruskal-Wallis test and Dunn's multiple comparison test (For 'Age', 'Duration from index presentation to Operation', 'Total bilirubin', 'ALT', 'AlkP' and 'GGT')

Among those who underwent a cholecystectomy (n=276, 44.6%), there was no difference in the time period from patient’s index hospital presentation to the day of operation between patient groups, even between those with (median: 3.0 days [2.0-4.8]) and without (median: 4.0 days [2.0-7.5]) complications of gallstones (P=0.12). Interestingly, there was no statistical difference between the two gallstone disease subgroups (uncomplicated: n=11, 17.2% vs. complicated: n=18, 15.7%; P=0.790) with regard to CT investigation, but there was a significantly different proportion of patients with complicated gallstone disease (n=43, 37.4%) who underwent MRCP compared to those with an uncomplicated disease (n=34, 53.1%) (P=0.041) (**Supplementary Table 1**). Serum ALT, AlkP and GGT activities were higher in patients with uncomplicated gallstone diseases compared to complicated diseases (Table 1).

### HMGB1

There was no significant difference in plasma HMGB1 concentration between patients with and without gallstone disease (*gallstone diseases*: 2.37 ng/mL [1.40-3.53]; *non-gallstones*: 2.31 ng/mL [1.22-3.60]; P=0.480) or between the four different study groups (P=0.584) (Table 2 and **Supplementary Figure 1**).

**Table 2.** Biomarker concentrations.

|  | Overall | NSAP<sup>1</sup> | Uncomplicated<sup>2</sup> | Complicated<sup>3</sup> | Others | P-value<sup>4</sup> |
| --- | --- | --- | --- | --- | --- | --- |
| miRNA-122 (x10<sup>4</sup> copies/ml) |  |  |  |  |  |  |
| Median | 1.81 | 0.90 | 5.72 | 2.26 | 1.13 | <0.001 |
| IQR | 0.54-7.0 | 0.41-2.23 | 0.97-16.85 | 0.75-7.86 | 0.42-4.42 |  |
| 95% CI | 6.21,16.40 | 4.32,26.42 | 9.51,22.24 | 1.54,22.82 | 2.26,8.23 |  |
| HMGB1 (ng/ml) |  |  |  |  |  |  |
| Median | 2.34 | 2.08 | 2.60 | 2.29 | 2.70 | 0.584 |
| IQR | 1.34-3.55 | 1.26-3.50 | 1.30-4.67 | 1.44-3.37 | 1.18-4.16 |  |
| 95% CI | 2.13,2.60 | 1.50,2.59 | 2.05,3.01 | 2.02,2.56 | 1.90,3.10 |  |
| cck-18 (U/L) |  |  |  |  |  |  |
| Median | 114.60 | 100.33 | 139.71 | 111.35 | 114.84 | 0.009 |
| IQR | 87.40-171.53 | 75.05-135.40 | 98.15-228.71 | 87.17-171.83 | 87.12-181.26 |  |
| 95% CI | 105.02,123.67 | 87.23,112.47 | 121.82,168.56 | 100.31,134.24 | 95.27,125.78 |  |
| flk-18 (U/L) |  |  |  |  |  |  |
| Median | 202.99 | 107.08 | 322.55 | 236.34 | 205.54 | <0.001 |
| IQR | 102.63-450.55 | 66.39-168.43 | 133.43-662.04 | 136.63-429.38 | 101.81-462.56 |  |
| 95% CI | 170.60,237.24 | 78.90,149.70 | 202.10, 479.32 | 184.91,304.06 | 151.78,325.69 |  |

<sup>1</sup> Non-Specific Abdominal Pain<sup>2</sup> Biliary colic, Aseptic Choledocholithiasis<sup>3</sup> Cholecystitis, Cholangitis, Gallstone Pancreatitis<sup>4</sup> Kruskal-Wallis test and Dunn's multiple comparison test

### miRNA-122

#### miRNA-122 in gallstone diseases and non-gallstone diseases

Circulating miRNA-122 concentration was higher in patients with gallstone disease than in those without gallstones (2.89 × 10^4^ copies/ml [0.88-9.94 × 10^4^] vs. 0.95 × 10^4^ copies/ml [0.41-3.30 × 10^4^]; P<0.001) (Figure 2A). ROC analysis (Figure 2B) of miRNA-122 for patients with and without gallstone disease produced an AUC of 0.66. The calculated PPV and NPV at 90% specificity were 72.5% and 39.8% respectively (Table 3).

**Figure 2A:**
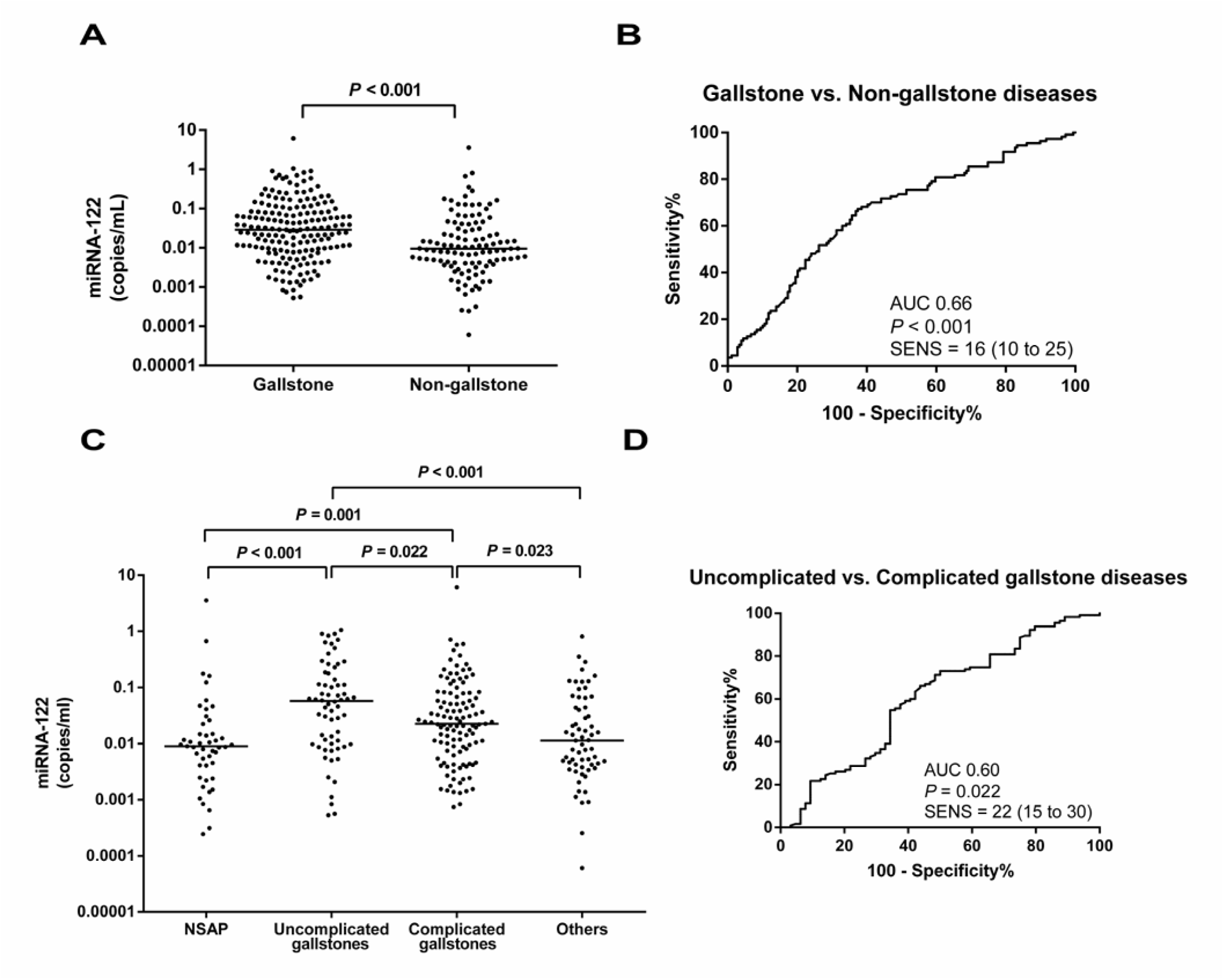
Plasma miRNA-122 values in gallstone and non-gallstone diseases normalised to miRNA-39. Each data point represents an individual. The horizontal line in each patient group represents the median value. Statistical significances (P value) by Kruskal Wallis test and Dunn’s multiple comparison test is shown in the figure for between group comparison. **B.** Receiver Operating Characteristic (ROC) curve of miRNA-122 for gallstone versus non-gallstone diseases. **C.** miRNA-122 values for the four main patient groups. **D.** ROC curve of miRNA-122 values for uncomplicated versus complicated gallstone diseases.

**Table 3.** Biomarker evaluation by receiver-operator characteristics.

|  | ROC AUC<br>(95% CI) | Sensitivity (%)<br>(95% CI) | PPV (%)<sup>1</sup> | NPV (%)<sup>2</sup> | P-value<sup>3</sup> |
| --- | --- | --- | --- | --- | --- |
| miRNA-122 |  |  |  |  |  |
| Gallstone vs.Non-gallstone<sup>4</sup> | 0.66(0.59-0.72) | 16(10-25) | 72.5 | 39.8 | <0.001 |
| Uncomplicated<sup>5</sup> vs. Complicated<sup>6</sup> | 0.60(0.51-0.69) | 22(15-30) | 56.0 | 67.5 | 0.022 |
| cck-18 |  |  |  |  |  |
| Gallstone vs.Non-gallstone<sup>4</sup> | 0.57(0.50-0.64) | 6(3-13) | 50.0 | 7.1 | 0.041 |
| Uncomplicated<sup>5</sup> vs. Complicated<sup>6</sup> | 0.59(0.50-0.68) | 14(8-22) | 45.0 | 65.4 | 0.049 |
| flk-18 |  |  |  |  |  |
| Gallstone vs. Non-gallstone<sup>4</sup> | 0.63(0.56-0.70) | 20(13-29) | 76.6 | 40.9 | <0.001 |
| Uncomplicated<sup>5</sup> vs. Complicated<sup>6</sup> | 0.56(0.47-0.65) | 12(7-20) | 42.1 | 65.0 | 0.160 |

<sup>1</sup> Positive predictive value<sup>2</sup> Negative predictive value<sup>3</sup> Kruskal-Wallis test and Dunn's multiple comparison test<sup>4</sup> NSAP and Other diseases<sup>5</sup> Biliary colic, Aseptic choledocholithiasis<sup>6</sup> Cholecystitis, Cholangitis, Gallstone pancreatitis

#### miRNA-122 in uncomplicated and complicated gallstone diseases

Patients with uncomplicated gallstone disease had a significantly higher miRNA-122 concentration than those with complicated gallstone disease (5.72 × 10^4^ copies/ml [0.97-16.85 × 10^4^] vs. 2.26 × 10^4^ copies/ml [0.75-7.86 × 10^4^]; P=0.022) (Table 2 and Figure 2C). ROC analysis (Figure 2D) comparing uncomplicated and complicated gallstone diseases yielded an AUC of 0.60 (Table 3) (**Supplementary Figure 2** and **Supplementary Table 2 and 5** illustrate the gallstone disease subgroups).

### cck-18

#### cck-18 in gallstone diseases and non-gallstone diseases

Plasma cck-18 concentration was significantly higher in patients with gallstone disease than those without (121.96 U/L [89.65-184.17]; vs. 104.57 U/L [84.23-150.55]; P=0.041) (Figure 3A). A ROC curve analysis of cck-18 in patients with and without gallstones yielded an AUC of 0.57 (Figure 3B), with a calculated PPV of 50.0% and NPV of 37.1% (Table 3).

**Figure 3A:**
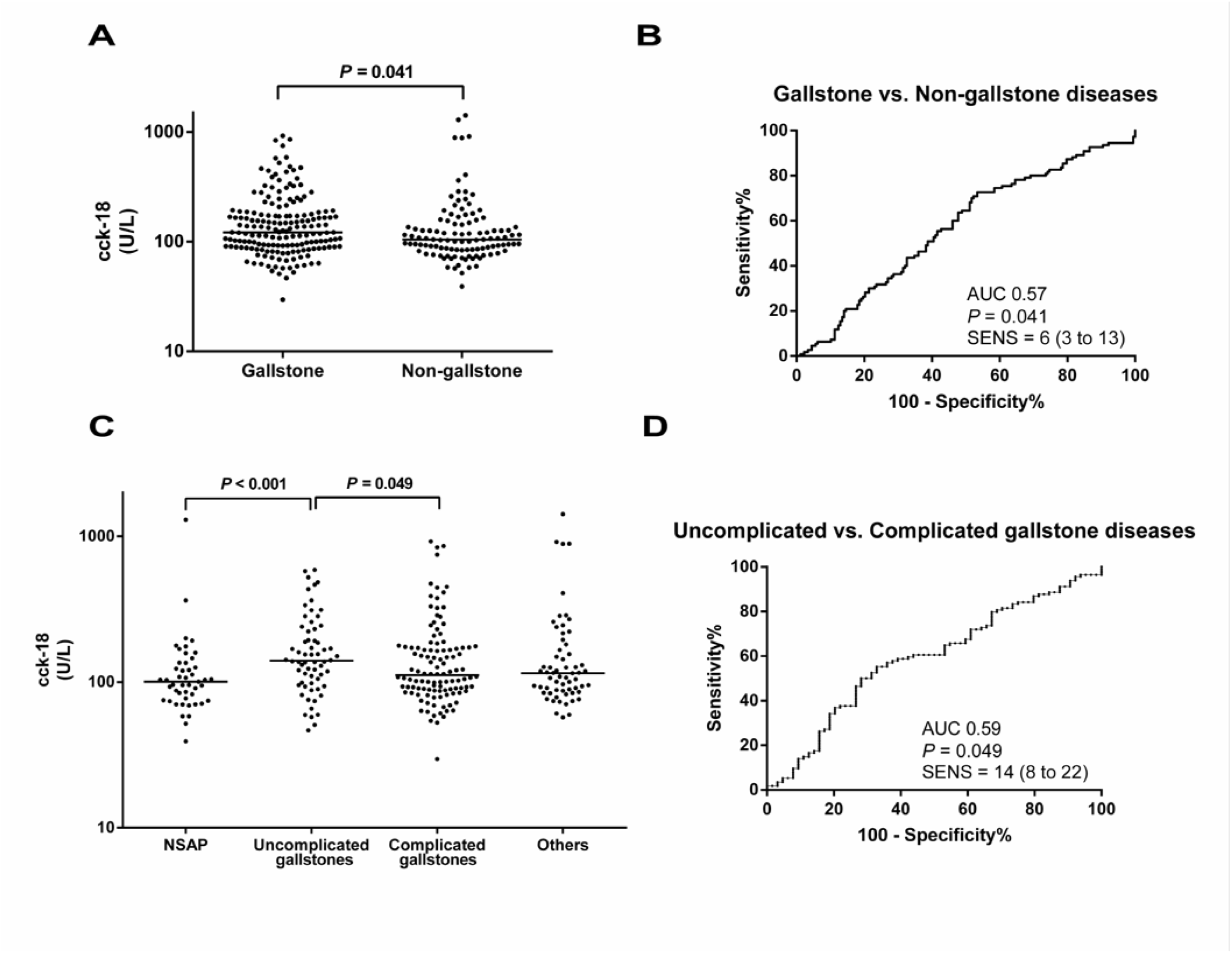
Plasma concentrations of cck-18 protein in gallstone and non-gallstone diseases. Each data point represents an individual. The horizontal line in each patient group represents the median value. Statistical significances (P value) Kruskal Wallis test and Dunn’s multiple comparison test is shown in the figure for between group comparison. **B.** ROC curve of cck-18 for gallstone versus non-gallstone diseases. **C.** cck-18 values for the four main patient groups. **D.** ROC curve of cck-18 values for uncomplicated versus complicated gallstone diseases.

#### cck-18 in uncomplicated and complicated gallstone diseases

cck-18 concentration was higher in patients with uncomplicated (139.71 U/L [98.15-228.71]) gallstone diseases than those with complications (111.35 U/L [87.17-171.83]) P=0.049) (Table 2 and Figure 3C). This translated to an AUC of 0.59 (Figure 3D) with PPV of 45.0% and NPV of 65.4% at 90% specificity (Table 3) (**Supplementary Figure 3** and **Supplementary Table 3** and **5** illustrate gallstone disease subgroups).

### flk-18

#### flk-18 in gallstone diseases and non-gallstone diseases

Similarly, flk-18 concentration was significantly higher in patients with gallstone disease than those without (252.38 U/L [136.63-525.67]; vs. 151.84 U/L [81.96-330.30]; P<0.001) (Figure 4A). A ROC curve analysis (Figure 4B) of flk-18 in patients with and without gallstones yielded an AUC of 0.63 (Table 3).

**Figure 4A:**
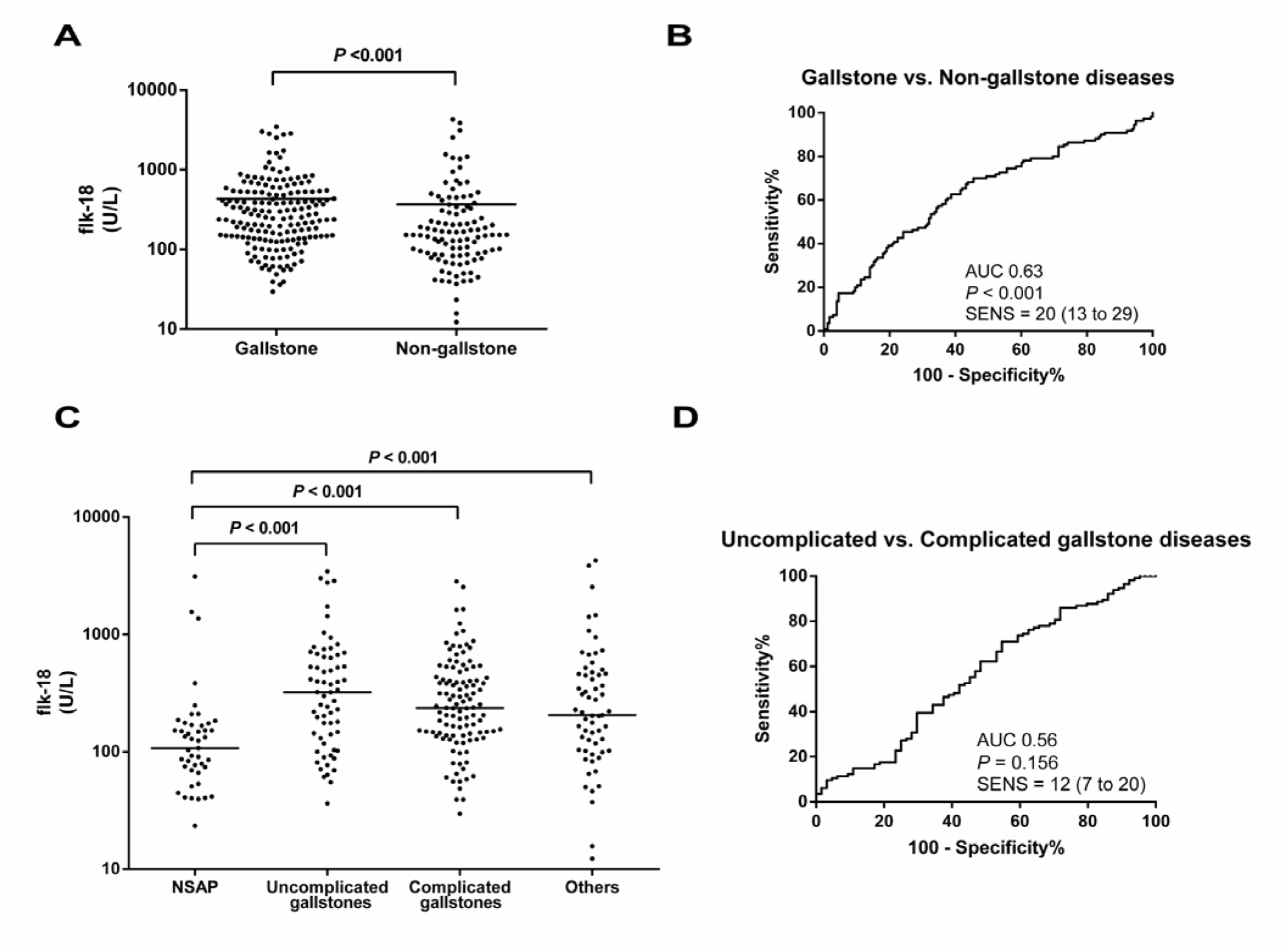
Plasma flk-18 protein concentrations in gallstone and non-gallstone diseases. Each data point represents an individual. The horizontal line in each patient group represents the median value. Statistical significances (P value) by Kruskal Wallis test and Dunn’s multiple comparison test isshown in the figure for between group comparison. **B.** ROC curve of flk-18 for gallstone versus non-gallstone diseases. **C.** flk-18 concentrations for the four main patient groups. **D.** ROC curve of flk-18 values for uncomplicated versus complicated gallstone diseases.

#### flk-18 in uncomplicated and complicated gallstone diseases

There was no significant difference in flk-18 concentration between the two subgroups of gallstone diseases (P=0.160) (Table 2 and Figure 4C) (**Supplementary Figure 4** and **Supplementary Table 4** and **5** illustrate gallstone disease subgroups).

## DISCUSSION

Early identification of individuals with potentially serious complications of gallstones from the denominator pool of all persons presenting with biliary-type symptoms is of translational importance. In this study, we report that plasma miRNA-122, cck-18 and flk-18 concentrations were significantly higher in patients with gallstone diseases than in non-gallstone diseases. However, the degree of overlap between groups was high, thus detracting from the utility of this approach to accurately stratify on an individual patient basis.

The biomarkers measured in this paper have robust evidence supporting their utility in preclinical models and the clinical setting for the early, sensitive and specific identification of hepatocellular injury following paracetamol overdose (an archetypal hepatocyte toxic agent) and also hypoxic hepatitis. Published papers have suggested this panel of biomarkers may also have utility with regard to acute cholestatic pathology induced by gall stones.

Our finding that plasma miRNA-122 concentration was increased in patients with gallstones agrees with that of Shifeng *et al*.^6^, who compared their disease cohort with ‘healthy’ control groups. However, their study yielded a much higher AUC on the ROC analysis, 0.93 with 77.4% sensitivity and 96.4%. The probable explanation for this is that our comparison group was patients admitted acutely under general surgery with a non-gallstone related diagnosis, who therefore may have had deranged inflammatory markers and/or LFTs due to another pathological process. This demonstrates the importance of clinically relevant control groups in biomarker discovery and validation studies.

In the present study, there was no significant difference in plasma HMGB1 concentration across all comparisons. This is consistent with a study by Shi *et al*.^7^which demonstrated HMGB1 over expression in gallbladder cancers but not in benign GB tissues or cholelithiasis.

Plasma cck-18 and flk-18 were significantly higher in patients with gallstones, in agreement with studies by others^14,22^ who examined gallbladder tissue, bile and serum. In our study, patients with cholecystitis were not further subdivided into active/inactive and acute/chronic subtypes due to sample sizing. Nevertheless, Simopoulos *et al.*^14^ demonstrated no difference in total k-18 and cck-18 between active and inactive chronic cholecystitis.

We recognize some important limitations in this study. This was a pilot study to determine whether there is a signal of biomarker utility in this context of use to take forward into larger, multi-centre studies. We believe that the results presented in this paper are clear in their failure to support utility with regard to patient stratification at hospital presentation. We did not proceed to perform multi-variable analysis given the near total overlap between the primary comparisons. Serial patient blood sampling for biomarker analyses may have also been useful to determine the true nature of biomarker concentration changes during the disease progress.

In conclusion, microRNA-122 and cleaved cytokeratin-18 plasma concentrations are elevated in patients with common bile duct complications of gallstones but are not sufficiently discriminatory to be progressed as clinically useful biomarkers in this context of use. Their clinical utility is in the context of acute hepatocyte injury.

## Study Contributors

*Edinburgh Emergency Surgery Study Group, Royal Infirmary of Edinburgh, NHS Lothian*: Graeme Couper, Christopher Deans, Gavin G. P. Browning, Anna M. Paisley, Bruce Tulloh, Richard J. E. Skipworth, Simon Paterson-Brown, Rajan Ravindran, Andrew de Beaux, Ijeoma Azodo.

## PREVIOUS COMMUNICATIONS

International Surgical Congress of the Association of Surgeons of Great Britain and Ireland (ASGBI) 2016, Belfast, Ireland. Oral presentation as Short Paper. Basic and Applied Clinical Science session, on 11^th^ May 2016.Presentation number 370. Abstract published in British Journal of Surgery: “The role of microRNA biomarkers to predict complications of gallstones”. [Aug 2016. Vol 103, Issue S6, Page 17]

## ACKNOWLEDGEMENTS

We are grateful for the expert assistance of the Edinburgh Clinical Research Facility and Dr. Kolamunnage Dona, Biomedical Statistician, University of Liverpool, UK. This project was funded by the Edinburgh and Lothians Health Foundation.

